# Ventrolateral prefrontal cortex in macaques guides decisions in different learning contexts

**DOI:** 10.1101/2024.09.18.613767

**Authors:** Atsushi Fujimoto, Catherine Elorette, Satoka H. Fujimoto, Lazar Fleysher, Brian E. Russ, Peter H. Rudebeck

## Abstract

Flexibly adjusting our behavioral strategies based on the environmental context is critical to maximize rewards. Ventrolateral prefrontal cortex (vlPFC) has been implicated in both learning and decision-making for probabilistic rewards, although how context influences these processes remains unclear. We collected functional neuroimaging data while rhesus macaques performed a probabilistic learning task in two contexts: one with novel and another with familiar visual stimuli. We found that activity in vlPFC encoded rewards irrespective of the context but encoded behavioral strategies that depend on reward outcome (win-stay/lose-shift) preferentially in novel contexts. Functional connectivity between vlPFC and anterior cingulate cortex varied with behavioral strategy in novel learning blocks. By contrast, connectivity between vlPFC and mediodorsal thalamus was highest when subjects repeated a prior choice. Furthermore, pharmacological D2-receptor blockade altered behavioral strategies during learning and resting-state vlPFC activity. Taken together, our results suggest that multiple vlPFC-linked circuits contribute to adaptive decision-making in different contexts.

## Introduction

In order to obtain the best possible outcome, organisms must flexibly adjust their behavior depending on environmental context^1^. For example, you have probably already learned through trial and error which transportation method, whether it be taxi, bus, or subway is best to take to get you to the airport in your hometown. However, you would need to reassess your strategy and learn which option would be most reliable to take when faced with the same task in an unfamiliar context like Tokyo or London. Such flexible adjustment of behavior depending on the context is vital for optimal behavior in an uncertain world. Its failure can be catastrophic, such as in gambling disorder, in which patients often lack cognitive flexibility and engage in risky behaviors despite significant economic losses^2–4^, or in schizophrenia, which is known to be associated with inflexible decisions based on delusional beliefs^5, 6^.

In non-human primates, the ventrolateral part of prefrontal cortex (vlPFC), especially Walker’s area 12, is critical for this type of learning and decision-making in uncertain environments. Specifically, fMRI and single neuron recording studies show that activity in vlPFC represents outcome probability and integrates this information into subjective value computations^7–11^. Studies that have either lesioned or inactivated vlPFC have reported that this area causally contributes to probabilistic learning and decision-making^8, 12^, but is not required when the associations between stimuli and rewards are deterministic^13^. A recent study also highlighted that dopaminergic projections to vlPFC and premotor cortex are critical for making choices in stochastic environments^14^ again indicating that a properly functioning vlPFC is essential for probabilistic decision-making.

In situations where people or animals have to adaptively determine the best course of action based on probabilistic feedback, they often use ‘win-stay/lose-shift’ behavioral strategies. Put simply, subjects pursue previously-rewarded choices and avoid previously-unrewarded choices^15^. Correctly applying such strategies to guide behavior in a given context can increase the rate of reward, and their use has been linked to the integrity of vlPFC^7, 8, 12^. A role for vlPFC in the use of such strategies, however, may run counter to the view that vlPFC is essential for learning from probabilistic feedback. One possibility is that the role of vlPFC in learning and decision-making varies depending on the learning context, that is, whether the stimulus-reward associations are known or must be learned. vlPFC may achieve this by differentially interacting with other cortical and subcortical areas, but the specific circuits are unclear^16–18^.

To test the role of vlPFC in learning and the use of strategies to guide behavior in different contexts, we first conducted a functional MRI experiment in awake macaque monkeys, while they participated in a probabilistic learning task. During the task, subjects completed blocks of trials under different learning contexts; either new stimulus-reward associations had to be learned (novel context) or knowledge about previously-learned associations could be used to guide choices (familiar context). In a second experiment, we examined the role of dopamine in influencing behavior across learning contexts through systemic injection of selective dopamine receptor antagonists. Finally, we then conducted anesthetized functional MRI under the same pharmacological challenge and specifically looked at the effect of dopaminergic modulation on vlPFC activity. Thus, this series of experiments allowed us to test how the role of vlPFC, and its interactions with cortical and subcortical areas, varies depending on the contextual modulation of learning and strategy use. The data indicate that vlPFC-linked pathways make distinct contributions to decision-making under different learning contexts.

## Results

### Animals exhibit context-dependent behavioral adaptation in a probabilistic learning task

Monkeys (N = 4) performed a probabilistic learning task for fluid rewards while they underwent whole-brain functional neuroimaging (**Fig. 1A-C**). On each trial, subjects chose between two visual stimuli presented on a monitor that were randomly selected from a larger pool of three stimuli. Each stimulus was associated with either 0.9, 0.5, and 0.3 probability of receiving a reward. Subjects completed trials in two different task contexts or blocks. In novel blocks, visual stimuli that subjects had never seen before were presented, whereas in familiar blocks stimuli that subjects had previously learned about were presented. Novel and familiar blocks were pseudorandomly intermingled in each session and neural and behavioral data were collected and analyzed in an event-related manner (**Fig. 1D**).

**Figure 1.**
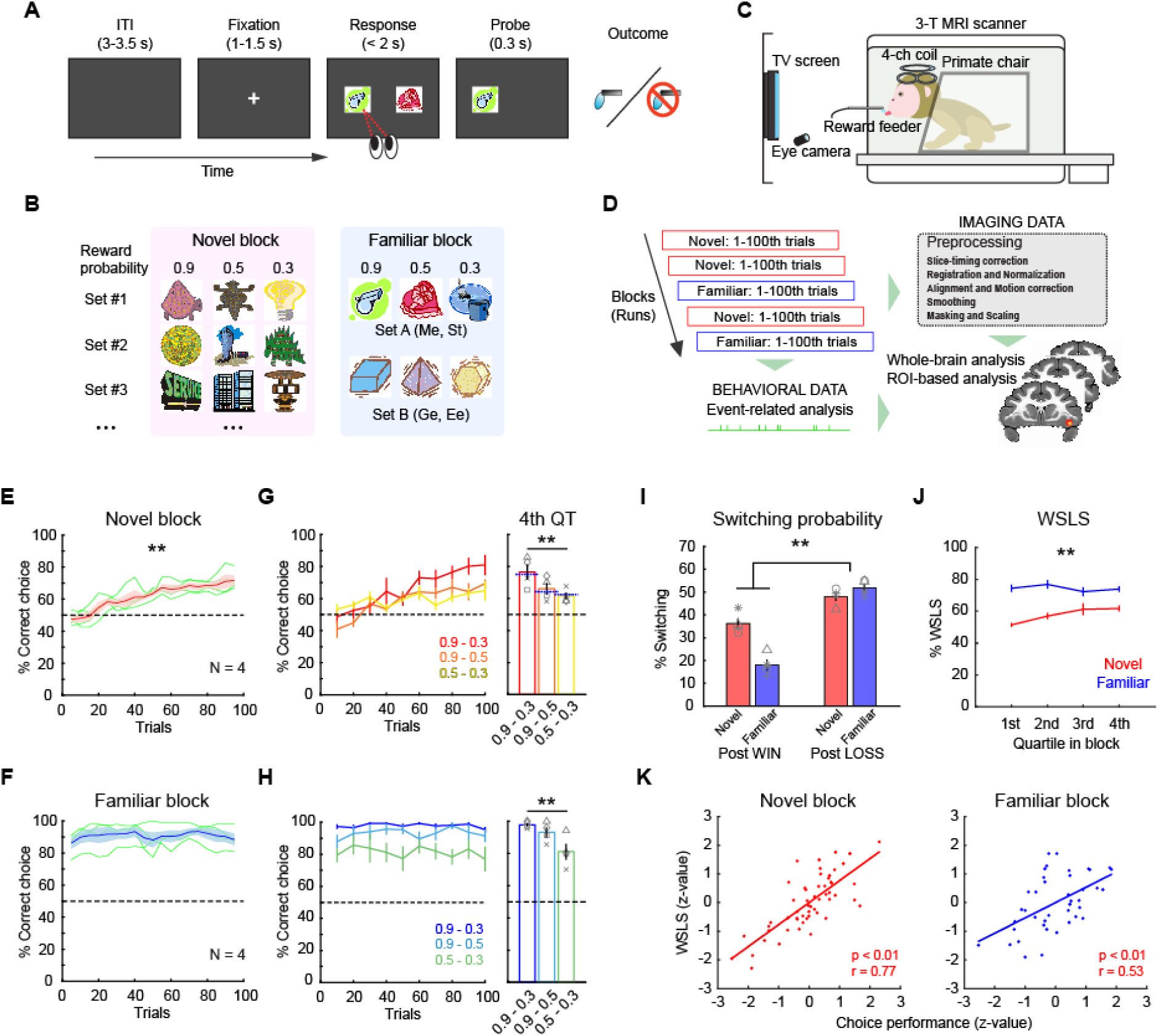
Probabilistic learning task and behaviors. (**A**) Trial sequence in a probabilistic learning task. On each trial, animals make a choice between two visual stimuli by eye movement to earn juice reward. (**B**) Stimulus sets in novel and familiar blocks. Each stimulus is associated with a reward probability of 0.9, 0.5, or 0.3. Different set of stimuli (Set A or B) are used by subject in familiar blocks. (**C**) Awake-fMRI setup. Subjects are placed in the sphynx position in the 3T MRI scanner in front of a display screen with an eye-tracking system, allowing them to perform tasks during functional scans. (**D**) Analysis pipeline. Neural and behavioral data are collected simultaneously and separately preprocessed offline for subsequent event-related analyses. (**E, F**) Choice performance in novel blocks (E) and familiar blocks (F). Average and SEM of choice performance (proportion of high-value option choice) of all monkeys (N = 4) are plotted. Asterisk indicates significant interaction of trial bin by block type (**p < 0.01, 2-way repeated-measures ANOVA). Dotted line indicates chance level. Green lines are individual performance. (**G, H**) Performance for each stimulus pair in novel blocks (G) and familiar blocks (H). Plots indicate performance in binned trials (left) where colors represent stimulus pair. Bar graph (right) indicates average performance for each stimulus pair in 4^th^ quartile. Asterisks indicate significant main effect of stimulus pair (**p < 0.01, 2-way repeated-measures ANOVA). Blue dotted line on the bar graph indicates the relative probability of a higher value option in each pair. Symbols represent individual animals. (**I**) Proportion of switching choices. Bars indicate average and SEM of switching probability for post-win trials and post-loss trials in novel and familiar blocks, respectively. Symbols represent each animal. Asterisk indicates significant interaction of block type by reward outcome (**p < 0.01, 2-way repeated-measures ANOVA). (**J**) Proportion of win-stay/lose-shift choices for novel (red) and familiar (blue) blocks in each quartile block (average and SEM). Asterisks indicate significant interaction of trial bin by block type (**p < 0.01, 2-way repeated-measures ANOVA). (**K**) Correlation between the proportion of WSLS and choice performance in novel (left) and familiar (right) blocks. Each dot represents individual blocks and lines indicate linear fitting of the data.

Subjects demonstrated distinct patterns of behavior across novel and familiar blocks. In novel blocks, subjects’ performance gradually improved as animals learned which option was associated with the highest probability of reward, whereas in familiar blocks performance was consistently at a high level (**Fig. 1E, F**). We split each block into equal 25-trial bins and found that performance in early trial bins was different between novel and familiar blocks, whereas later bins were not (2-way repeated-measures ANOVA, interaction of block type by trial bin, p < 0.01, F_(1,178)_ = 18.1). This indicates that the animals successfully learned new stimulus-reward associations in novel blocks while they maintained high performance across familiar blocks. Within both learning contexts (i.e., novel and familiar blocks) we found that correct performance in later bins reflected the relative reward probability associated with stimuli available on each trial (2-way repeated-measures ANOVA, main effect of stimulus pair, p < 0.01, F_(2,267)_ = 12.6) (**Fig. 1G, H**). This meant that the macaques were not always choosing the available option with the highest probability of reward but were distributing the frequency of their choices to match the relative option value. The response time (RT) also reflected the relative reward probability at each stimulus pair in both block types (2-way repeated-measures ANOVA, main effect of stimulus pair, p = 0.016, F_(2,267)_ = 4.2) (**Supplementary Fig. 1**). Such a pattern of responding is consistent with matching, a behavior whereby subjects distribute their responding to the available options^19, 20^.

Given that subjects exhibited aspects of matching behavior, which takes into account the outcome of the previous trial, we next looked at subjects’ use of reward delivery-based behavioral strategies. Here we found that animals demonstrated distinct behavioral strategies depending on the learning context. Overall, they tended to switch their choices more frequently following a ‘loss’ (unrewarded) trial compared to a ‘win’ (rewarded) trial, manifesting a win-stay/lose-shift (WSLS) pattern (**Fig. 1I**). This tendency was more pronounced in familiar blocks than novel blocks (2-way repeated-measures ANOVA, interaction of outcome by block type, F_(1,178)_ = 47.8, p < 0.01), indicating that the monkeys were more likely to apply WSLS strategy when learning was not required. Specifically, the proportion of WSLS trials was at chance in the early phase of novel blocks, but gradually increased toward the end of blocks, while it was maintained at a high level throughout familiar blocks (2-way repeated-measures ANOVA, interaction of trial bin by block type, F_(3,356)_ = 4.0, p < 0.01) (**Fig. 1J**). In addition, the proportion of WSLS trials was positively correlated with choice performance in both novel and familiar blocks (Pearson’s correlation, n = 55 and 42 for novel and familiar blocks, respectively, p < 0.01) (**Fig. 1K**), suggesting that the use of these strategies based on the learning context was advantageous for task performance. Taken together, these analyses show that the subjects adaptively used behavioral strategies to improve their task performance across the different learning contexts.

### Whole-brain encoding of outcome and learning context

Our behavioral analysis demonstrated that subjects adjusted their behavioral strategies between learning contexts, altering their decisions to stay or shift from previous choices depending upon the outcome. Consequently, we next set out to determine the network of brain areas that exhibited neural activity associated with the task. First, we analyzed whole brain functional neuroimaging data collected from subjects while they performed the task (**Fig. 1C, D**), looking for signals that were modulated either by learning context (**Fig. 2A**) or reward outcome (**Fig. 2B**). These analyses revealed that bilateral lateral and ventral frontal areas as well as the ventral temporal lobes were more active in the novel versus familiar blocks, while medial frontal areas showed greater activity in the familiar blocks (**Fig. 2A**). By contrast, reward receipt was associated with increased activity in ventrolateral frontal cortex as well as parts of sensorimotor cortex and ventral striatum, and decreased activity in dorsolateral prefrontal cortex (**Fig. 2B**).

**Figure 2.**
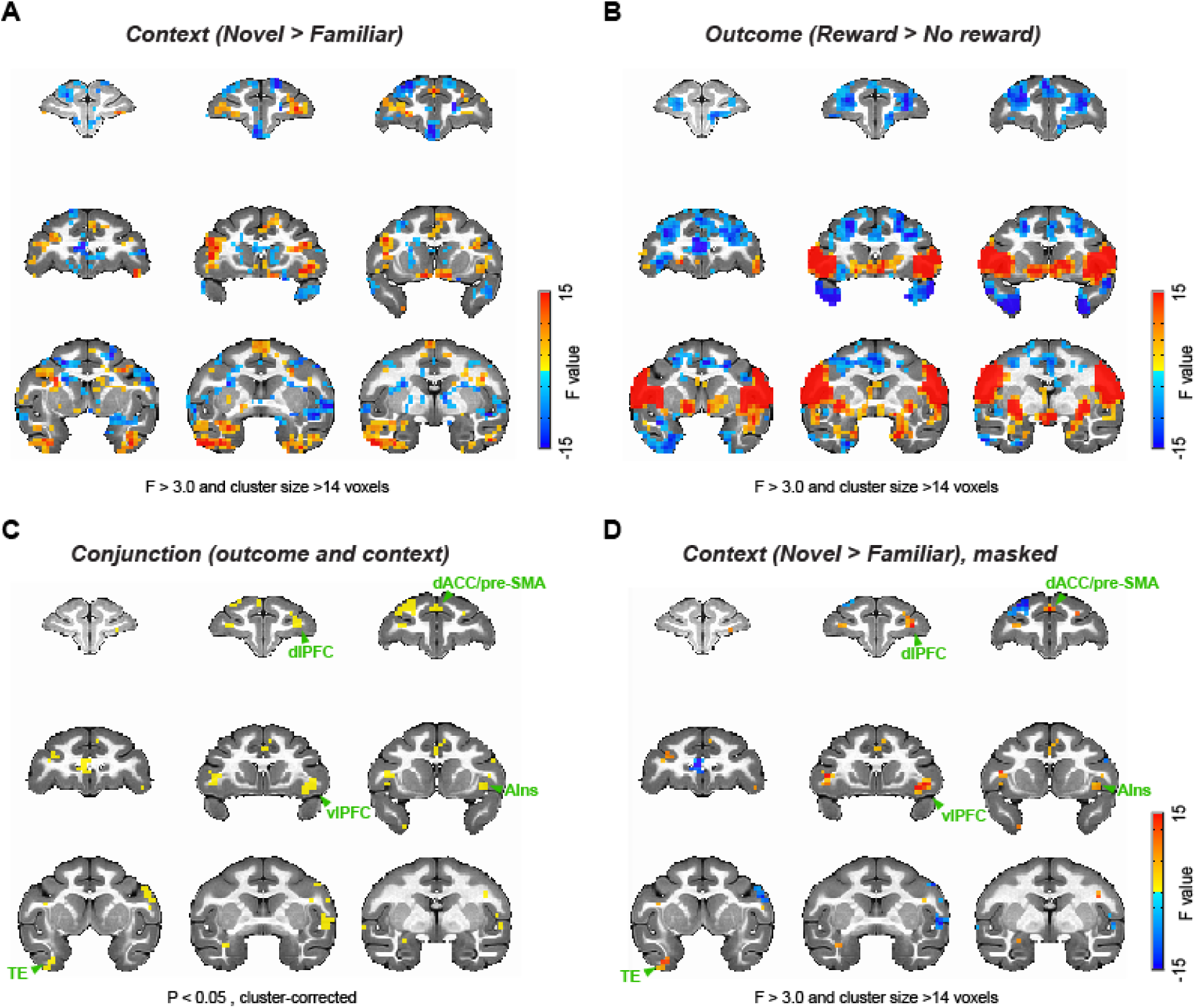
Whole-brain representations of learning context and outcome. (**A**) Whole-brain representations of learning context. Coronal slices (2.5 mm apart) are shown from anterior (top left) to posterior (bottom right) planes. Thresholded F-stat maps (p < 0.05, cluster-corrected) are superimposed on a standard anatomical template. Positive and negative F-stats (warmer and cooler colors) indicate more activity in novel blocks and in familiar blocks, respectively. (**B**) Whole-brain representations of reward outcome. Larger F-stats indicate more activity in rewarded than no reward trials. Data are displayed in the same manner as (A). (**C**) Conjunction analysis result. Clusters highlighted (yellow) significantly encoded both learning context (novel vs. familiar) and reward outcome (rewarded vs. no reward) at cluster-level correction (p < 0.05). (**D**) F-stats map of context coding (novel vs. familiar; A) masked for the clusters identified in the conjunction analysis (C). dlPFC: dorsolateral prefrontal cortex, dACC: dorsal anterior cingulate cortex, pre-SMA: pre-supplementary motor area, vlPFC: ventrolateral prefrontal cortex, AIns: anterior insula, TE: inferior temporal cortex.

Next, we conducted a conjunction analysis looking for areas showing activations that varied based on context and reward outcome. This analysis revealed clusters that encoded both context and reward outcome, in a distinct network of areas including vlPFC, dorsal anterior cingulate cortex (dACC), dorsolateral PFC, supplemental motor area (SMA), and inferior temporal cortex (TE) (2-way ANOVA, main effect of block type or outcome, p < 0.05 with cluster-correction, **Fig. 2C**). We then projected the effect of the novel versus familiar comparison back onto the areas that showed an interaction between the effects of context and reward outcome to visualize the strength of context encoding (**Fig. 2D**). This analysis showed that the vlPFC and dACC, the areas previously highlighted based on their potential role in probabilistic learning^18, 21^, are indeed associated with learning context and reward, and respond more strongly in the novel learning context compared to the familiar context. Taken together, these whole-brain analyses suggest that the activity in vlPFC varies based on the ongoing learning context and the preceding outcome. A full table of statistically significant clusters for this analysis can be found in **Supplementary Table 1**.

### vlPFC activity tracks outcome and behavioral strategy

Previous work has emphasized the critical role of neural activity in vlPFC in probabilistic learning^7, 8, 12^. Here we found that across all subjects, activity in bilateral vlPFC consistently varied with learning context (novel > familiar) and this effect was most clearly differentiated in the right vlPFC (**Fig. 2** and **Supplementary Fig. 2**). Consequently, we chose right vlPFC as a region of interest (ROI) for further analyses (**Fig. 3A**). In each novel and familiar block, the signal in vlPFC varied depending on the trial-by-trial outcome and whether subjects subsequently stayed with their prior choice or shifted to a different option (**Fig. 3B**). The average signal in this area around reward delivery (0-4 s after outcome) was marginally and negatively correlated with task performance in novel blocks (Pearson’s correlation, n = 51, p = 0.056), while there was no consistent relationship between performance and vlPFC activity in familiar blocks (n = 42, p = 0.57) (**Fig. 3C**). Such a pattern of effects potentially indicates that activity in vlPFC is higher during explorative behavioral adaptation when the macaques are learning new stimulus-reward associations.

**Figure 3.**
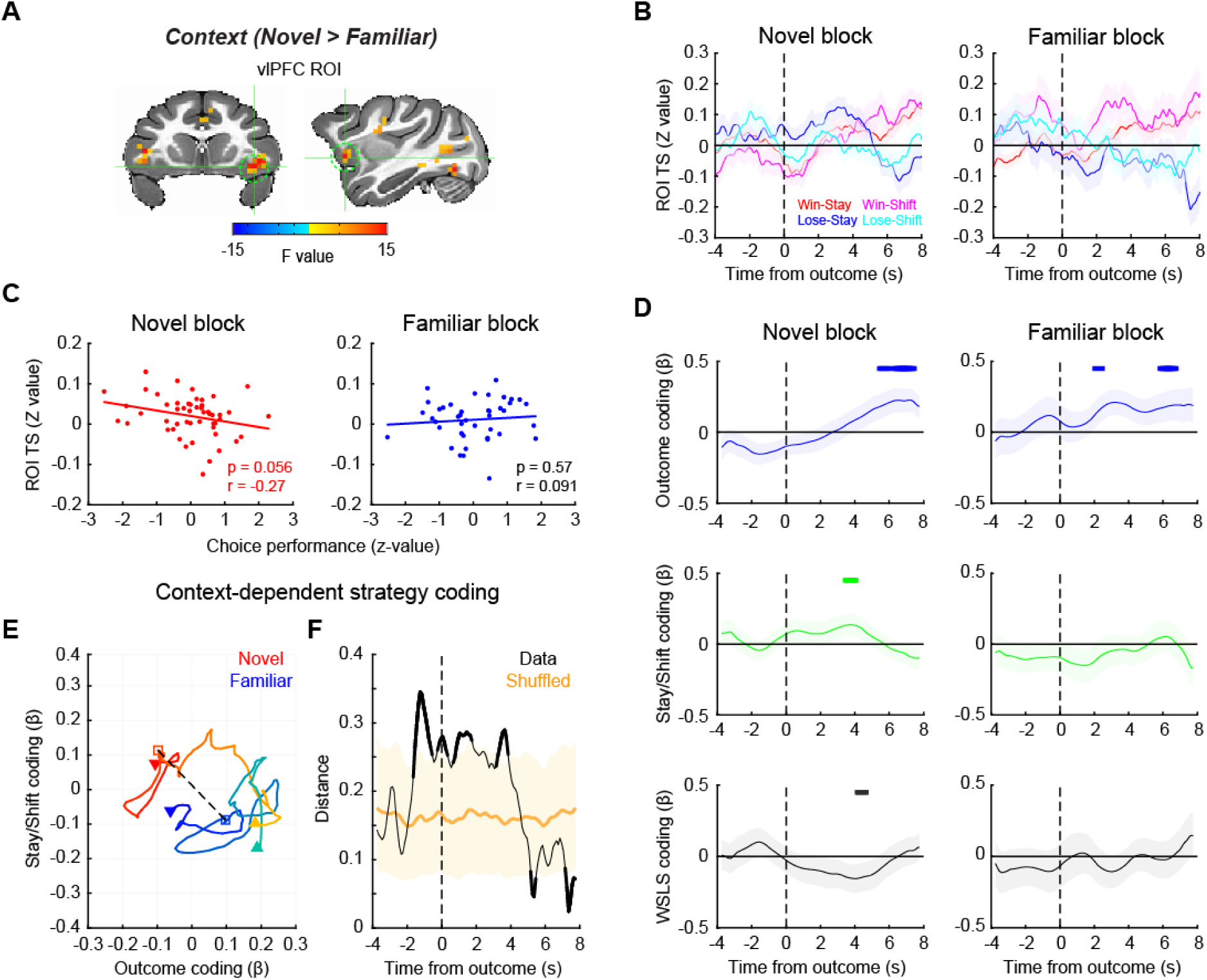
vlPFC signal encodes behavioral strategy during learning. (**A**) vlPFC ROI for time-series analysis. The map of F-stats of context coding (novel vs. familiar) are shown on coronal (left) and sagittal (right) planes of a standard anatomical template. A spherical ROI i defined based on the peak coordinates of context coding in the right vlPFC cluster. (**B**) ROI time-series around the outcome timing during novel (left) and familiar (right) blocks. Average and SEM of ROI time-series are plotted for win-stay, win-shift, lose-stay, lose-shift trials, respectively. (**C**) Correlation between vlPFC activity and choice performance in novel (left) and familiar (right) blocks. Each dot indicates a block and the line indicates a linear fitting of the scatter plot. (**D**) Regression analysis result. Beta coefficients for outcome coding (top), stay/shift decision coding (middle), and the interaction of outcome by stay/shift decision (i.e., WSLS behavioral strategy) coding (bottom) were computed using a sliding window analysis. The time-course of the beta coefficients were plotted around the timing of outcome (vertical dotted line) for each of novel (left) and familiar (right) blocks. Thick lines on the top of each panel indicate significant encoding compared to zero (p < 0.05 at 3 consecutive bins, rank-sum test). (**E, F**) Multidimensional analysis result. (**E**) Beta coefficients for outcome and stay/shift decision coding are plotted at each time point of novel (warmer colors) and familiar (cooler colors) block, with the passage of time represented as a gradient of colors. The dotted line and squares indicate the timing of outcome, and downward arrow and upward arrow indicate the start and end of the analysis window (from -4 to 8 seconds after the outcome), respectively. (**F**) The Euclidian distance between novel and familiar blocks was computed at each time point and plotted against the time. The shaded area indicates the 95% confidence interval of the shuffled data. The data that exceeded the 95% CI are represented by thick lines.

To more formally assess this relationship, we compared activity in vlPFC between learning contexts. A multiple-regression analysis was performed on the ROI time-series for each block type. To investigate the effects of different factors on signals in vlPFC, this analysis included the following factors as regressors: reward outcome of the present trial, stay/shift decision in a subsequent trial, and the interaction between these two, i.e. whether subjects were using a win-stay/lose-shift (WSLS) strategy. This analysis revealed that vlPFC encoded whether reward was delivered or not in a similar manner across both novel and familiar blocks (**Fig. 3D, top panels**). By contrast, only activity within vlPFC during novel blocks was related to subjects’ decision to stay or shift and their use of WSLS strategies (compare left and right side of **Fig. 3D, middle and bottom panels**). Thus, when subjects are actively learning stimulus-reward associations in novel blocks, activity within vlPFC is driven by both reward delivery as well as the behavioral strategy that the subjects are using.

To explore the time course of the differences in vlPFC signal between the novel and familiar blocks, we conducted a multidimensional analysis of activity related to reward outcome and whether subjects chose to stay or shift their behavior. To do this we projected the regression coefficients (beta values) from these two variables from 4 seconds before to 8 seconds after the outcome onto a 2-D space for each block type (**Fig. 3E**). We then measured the Euclidean distance between the projected beta values from the novel and familiar blocks at each time point, as a proxy of neural representational difference between the two contexts, that were plotted against the time relative to the outcome (**Fig. 3F**). This analysis revealed that the neural encoding of reward outcome and decisions to stay or shift in vlPFC in the two contexts most prominently diverged around the timing of the outcome (permutation test, p < 0.05). This indicates that the activity within vlPFC diverges at the timing when animals are adjusting their use of behavioral strategies depending on the context.

### vlPFC-ACC functional connectivity encoded behavioral strategy during learning

vlPFC is a hub of the frontal attention network and the salience network^16, 17^. Therefore, we next asked how functional networks centered on the vlPFC are associated with reward outcome and learning context using a generalized psycho-physiological interaction (gPPI) analysis^22^. We first mapped out voxels in the brain whose time-series showed interaction with the vlPFC seed time-series and reward outcome or context. Based on this, we then identified significant clusters that were modulated by the context that subjects were in (2-way ANOVA, main effect of block type, p < 0.05 with cluster-correction) (**Supplementary Fig. 3**). This analysis showed that the functional connections between vlPFC and ACC, mediodorsal thalamus (MD), dlPFC, and pre-motor areas were modulated by the learning context. A full table of statistically significant clusters is in **Supplementary Table 2**.

Among these, we first focused on the vlPFC-ACC functional connection (**Fig. 4A**), as both vlPFC and ACC are implicated in adaptive behavior^18^ and are known to be anatomically and functionally connected^16^. Here we found that vlPFC-ACC functional connectivity (FC) varied with the behavioral strategies used by the subjects after reward feedback (**Fig. 4B**). Specifically, FC between vlPFC and ACC increased around the time of outcome (rank-sum test, p < 0.05 at 3 consecutive bins) when the animals received reward and repeated the same choice (i.e., win-stay) or when the animals received no reward and subsequently changed their choice (i.e., lose-shift), representing a WSLS pattern in novel blocks (**Fig. 4B, left panels**). By contrast, in familiar blocks, the FC changes did not follow the WSLS pattern although some significant modulation was observed mainly before outcome period (**Fig. 4B, right panels**). Three-way ANOVA confirmed these effects as there was a significant interaction of block type by stay/shift decision and reward outcome, indicating that FC changes reflected WSLS pattern exclusively in novel blocks (**Supplementary Fig. 4**; F_(1,368)_ = 4.0, p = 0.046). This pattern of effects indicates that such connectivity was related to task performance, and indeed the functional connectivity between vlPFC and ACC in loss trials was marginally and negatively correlated with task performance in the novel context (Pearson’s correlation, n = 51, r = -0.26, p = 0.056). No such correlation was observed in the familiar blocks (n = 42, r = -0.061, p = 0.70) or for performance and FC on win trials (p > 0.30) (**Fig. 4C**). A sliding-window regression analysis showed that the FC between vlPFC and ACC was associated with the WSLS strategy around the time of outcome in novel blocks (permutation test, p < 0.05), while they were anti-correlated around the reward timing in the familiar context (**Fig. 4D**). This further suggests that the functional interaction between vlPFC and ACC is related to the context-dependent use of behavioral strategies.

**Figure 4.**
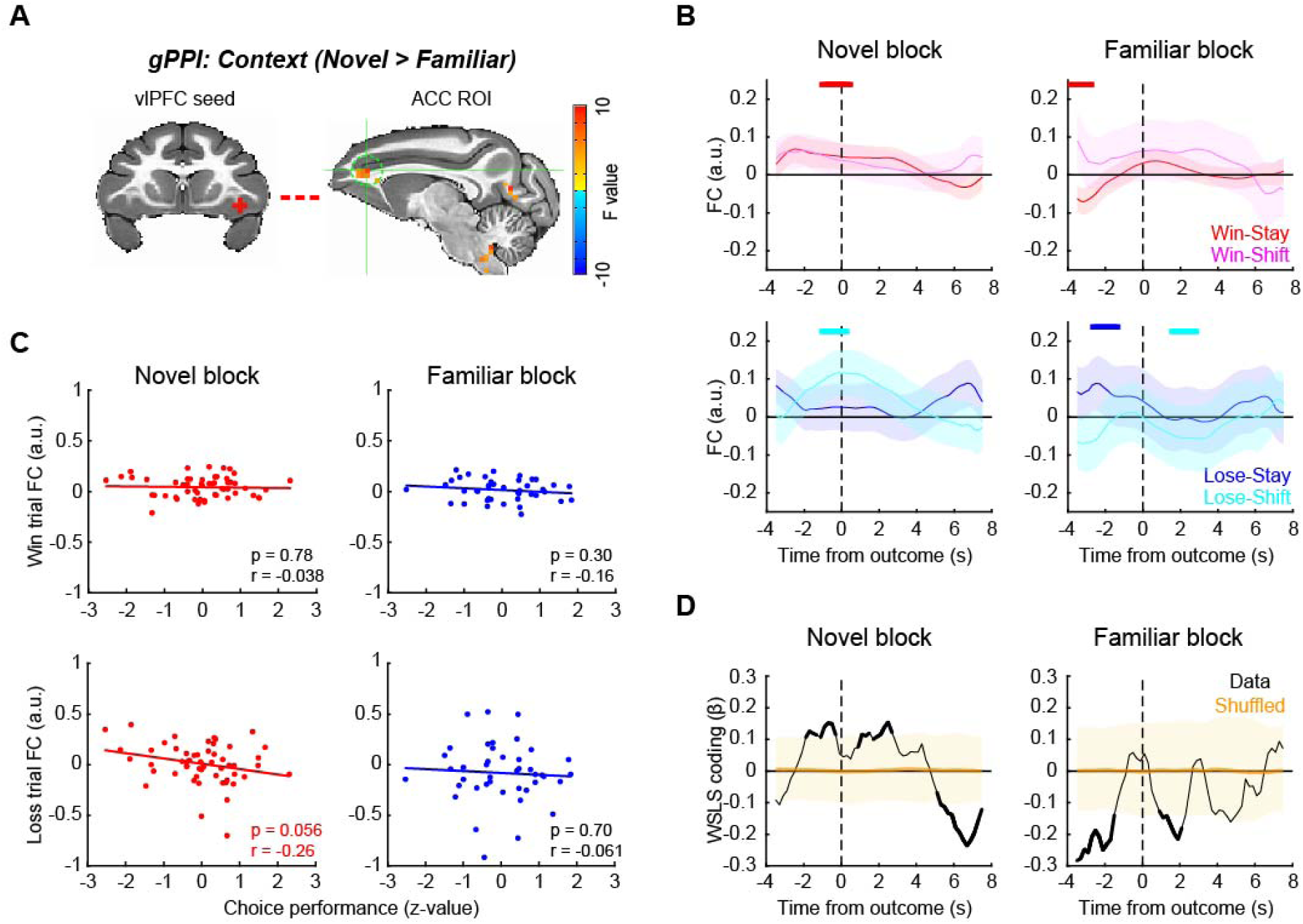
vlPFC-ACC functional connection encodes behavioral strategy during learning. (**A**) vlPFC seed and ACC ROI for functional connectivity analysis. ACC ROI (sagittal plane on the right) was defined based on generalized PPI analysis using right vlPFC seed. (**B**) FC time course around the outcome timing. vlPFC-ACC FC during novel (left) and familiar (right) blocks were computed using sliding window analysis and visualized for win-stay and win-shift trials (top) and lose-stay and lose-shift trials (bottom) separately. The plots are made around the outcome timing (vertical dotted lines). The thick lines on the top of each panel indicate significant FC compared to zero for color matched trials (p < 0.05 with rank-sum test at 3 consecutive bins). (**C**) Correlation between vlPFC-ACC FC and choice performance. The correlations were computed for win (rewarded) trials (top) and loss (unrewarded) trials (bottom), and for novel (left) and familiar (right) blocks separately. Each dot represents each block, and the lines are linear fitted to the data. (**D**) Time course of WSLS coding around the outcome. WSLS coding was computed as the interaction of outcome by stay/shift decision coding in a sliding window multiple-regression analysis. Shaded areas (yellow) indicate 95% confidence interval of the data, and the thick black lines indicate the significance of the data.

### vlPFC-MD functional connectivity reflects decision to stay with a choice during learning

Previous studies have shown that the fronto-thalamo pathway plays a critical role in learning and decision-making^23–25^. Consequently, we next focused on the functional connection between vlPFC and MD (**Fig. 5A**). The functional connectivity between vlPFC and MD specifically increased in trials that were followed by the repetition of the previous choice (i.e., stay decision) regardless of whether reward was delivered or not in novel blocks (**Fig. 5B, left panels**). This pattern was less pronounced in familiar blocks (**Fig. 5B, right panels** and **Supplementary Fig. 5**; three-way ANOVA, interaction of block type by stay/shift decision, F_(1,368)_ = 4.4, p = 0.036). Interestingly, the choice signal was not correlated to task performance in either block (**Fig. 5C**; p > 0.44), and behavioral strategy coding was primarily observed in familiar blocks (**Fig. 5D**). This result suggests that this functional connection between vlPFC and MD encodes execution of the decision to stay or switch *per se* that is not directly linked to the correct performance.

**Figure 5.**
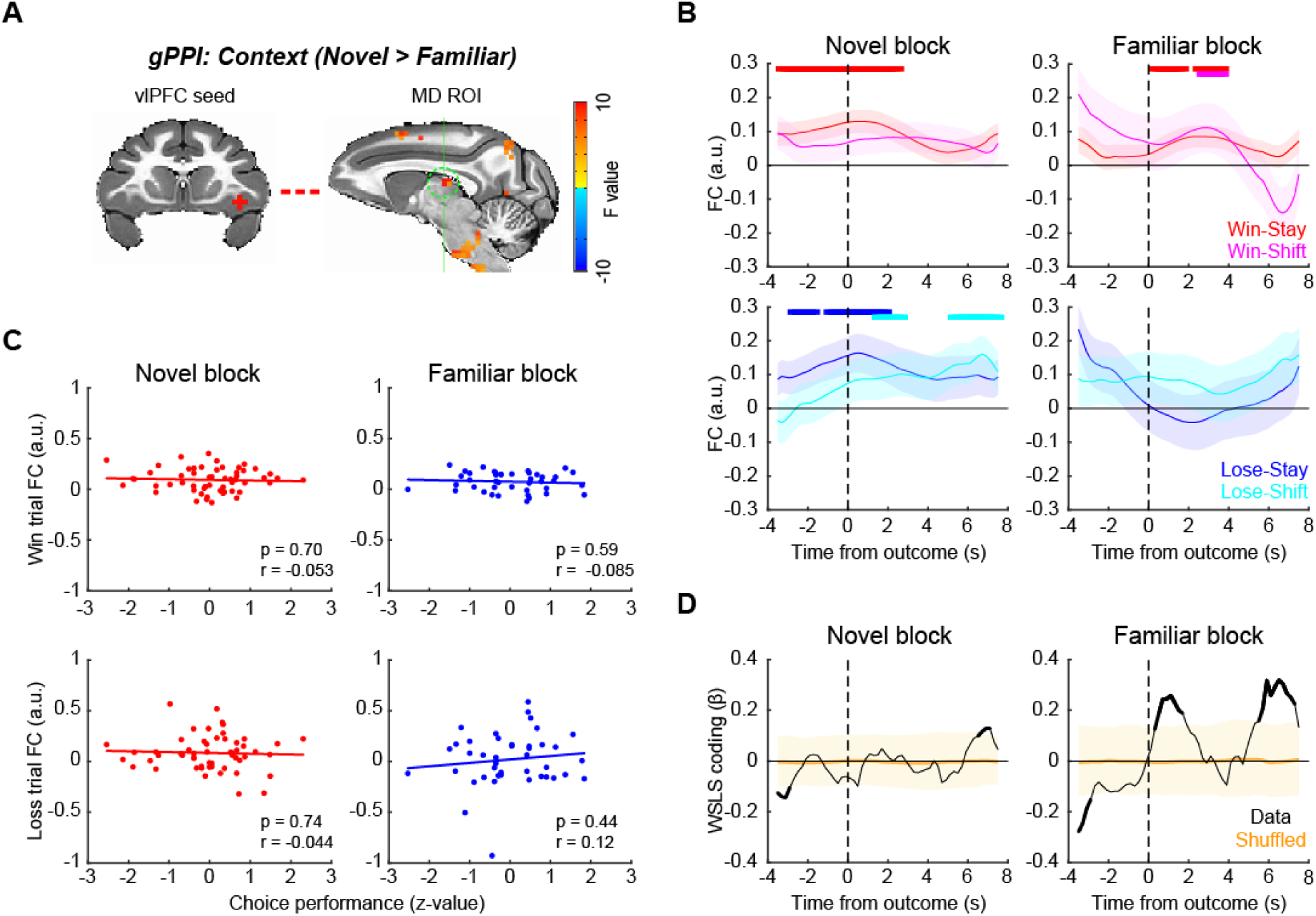
vlPFC-MD functional connection encodes decision to stay during learning. (**A**) vlPFC seed and MD thalamus ROI for functional connectivity analysis. (**B**) FC time course around the outcome timing. vlPFC-MD FC during novel (left) and familiar (right) blocks were computed for win-stay, win-shift, lose-stay, and lose-shift trials separately. The thick lines on the top of each panel indicate significant FC compared to zero for color matched trials (p < 0.05 with rank-sum test at 3 consecutive bins). (**C**) Correlation between vlPFC-MD FC and choice performance, for win trials (top) and loss trials (bottom) separately. Dots and lines indicate blocks and linear fitted line, respectively. (**D**) Time course of WSLS coding around the outcome. Shaded areas (yellow) indicate 95% confidence interval of the data, and the thick black line indicate the significance of the data.

We additionally looked at other vlPFC functional connections that showed significant novel vs familiar coding in the gPPI analysis (**Supplementary Fig. 3**). The functional connection between vlPFC and supplementary motor area (SMA) increased in the ‘stay’ trials during novel blocks in a pattern similar to that was observed with vlPFC-MD FC, although the interaction of block type by stay/shift decision was not significant (p = 0.30). The functional connection between vlPFC and dlPFC also showed changes depending on stay/shift decision in both block types, but again there was no significant interaction of block type by decision (p = 0.51). A lack of clear relationship between connectivity in these pathways and the learning context suggests that they might be more associated with different aspects of the learning context such as attention.

### Pharmacological manipulation of dopamine receptors affects vlPFC-mediated behavior

The prior analyses indicate that the WSLS strategy is crucial for adaptive behavior depending on the learning context, and this is in line with previous work showing that vlPFC plays a central role in probabilistic learning and decision-making^12, 14^. The release of dopamine in frontal cortex has been implicated in a variety of cognitive functions relevant to probabilistic learning, such a attention and working memory^26, 27^. Therefore, it is possible that the changes in vlPFC activity that were associated with behavioral strategy are mediated by the action of dopamine through cortical or subcortical dopamine receptors.

To address this, we conducted a pharmacological experiment with selective dopamine receptor antagonists SCH-23390 (D1 antagonist) and haloperidol (D2 antagonist) and assessed their effects on the performance of monkeys in the probabilistic learning task. The subject cohort in this experiment (N = 4) partially overlapped with the one used in our awake fMRI experiment (see **Table 1**), and the effects of the drugs on the proportion of correct choice and reaction times were analyzed in our previous paper in relation to resting-state functional connectivity^28^. Here, we specifically focused on the effects of systemically administered dopaminergic drugs on WSLS behaviors. D2 antagonist haloperidol, but not D1 antagonist SCH-23390 or saline, increased the proportion of WSLS responses preferentially in novel blocks (**Fig. 6A, B**). A two-way repeated-measures ANOVA demonstrated a significant main effect of drug dose in novel blocks but not in familiar blocks after haloperidol administration (novel: F_(2,440)_ = 3.9, p = 0.021; familiar: F_(2,396)_ = 0.072, p = 0.93), while there was no significant effect following SCH-23390 administration in either block type (p > 0.46). This result suggests that dopamine D2 receptors play a key role in modulating behavioral strategy during learning.

**Table 1.**
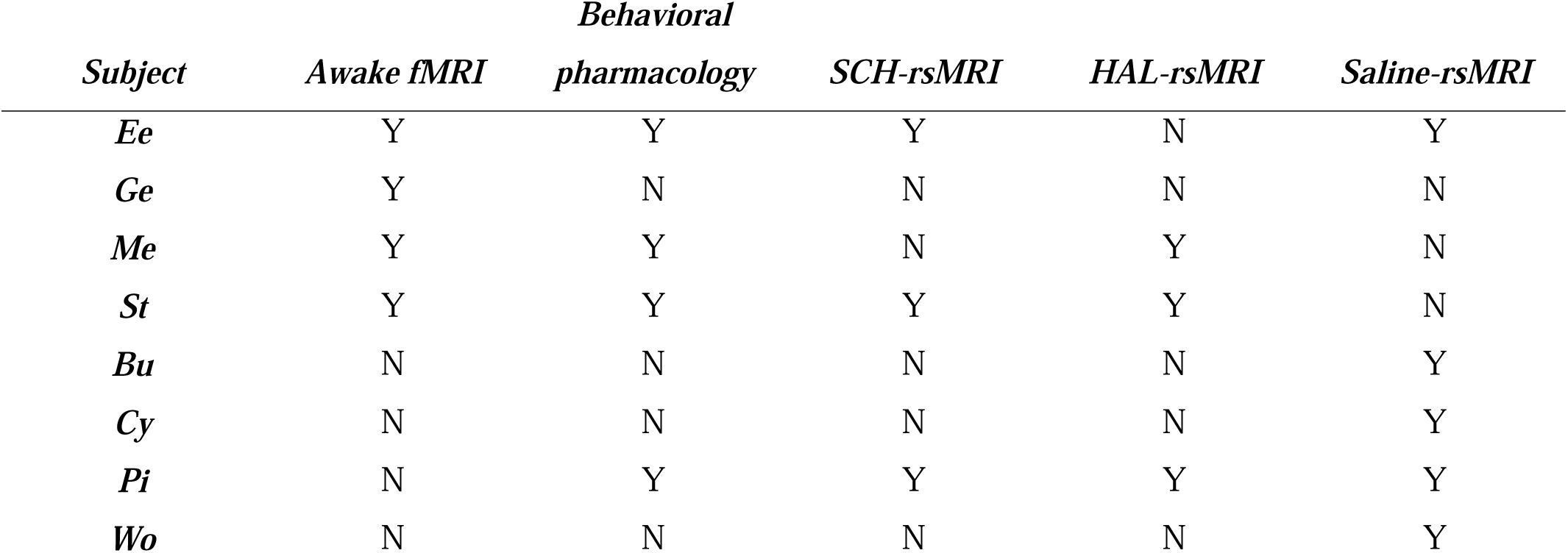
Assignment of experiments for each subject. Y and N indicate the condition that the data was collected and not collected, respectively. SCH: SCH-23390 (10 µg/kg), HAL: haloperidol (50 µg/kg). rsMRI: resting-state fMRI. Note that animals assigned to behavioral pharmacology experiments went through all drug treatment conditions.

**Figure 6.**
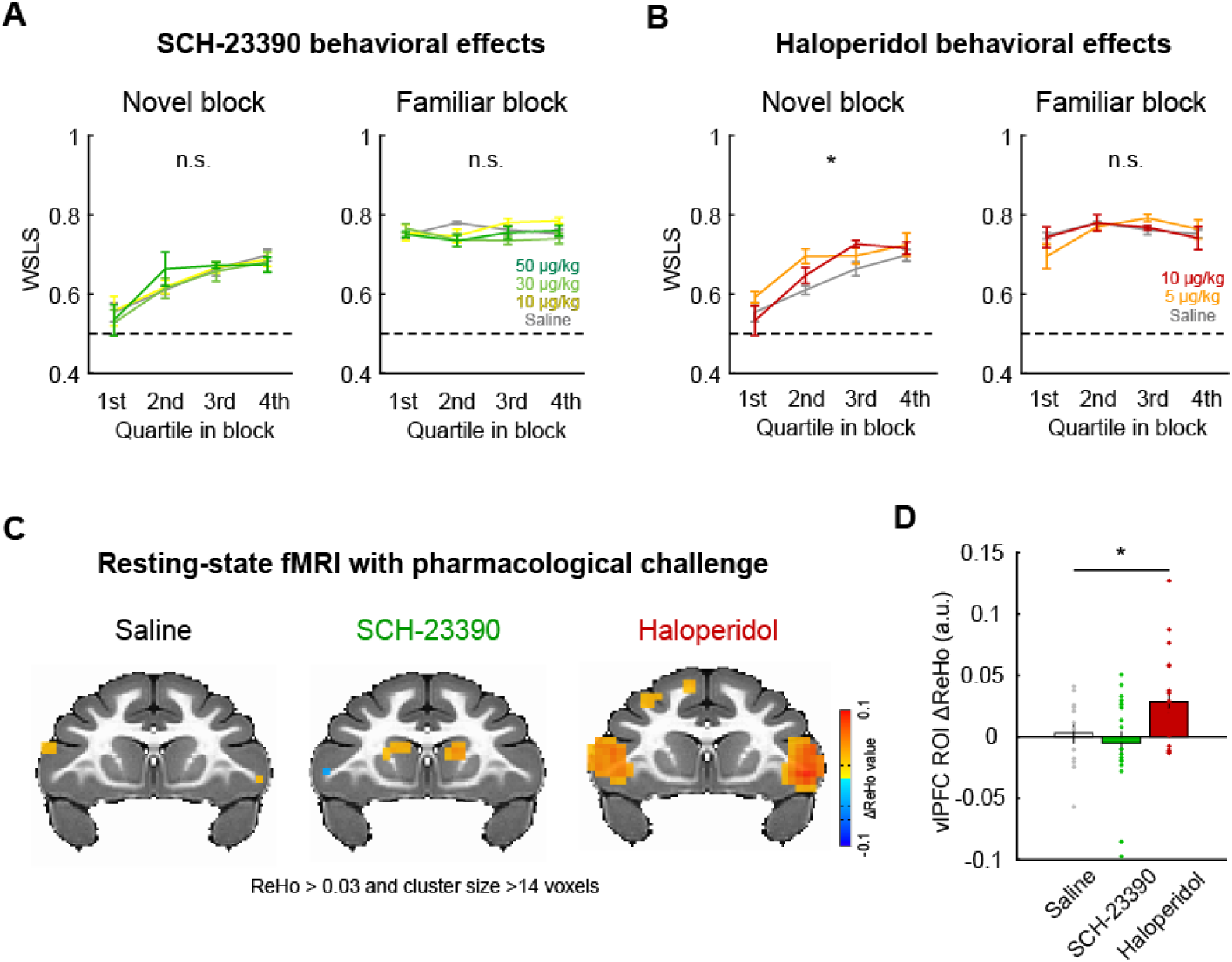
D2 receptor blocker enhanced vlPFC activity and promoted adaptive behavior. (**A**) The effect of D1 receptor antagonism on WSLS behavior. The proportion of WSLS trials in quartile blocks (average and SEM) are plotted for each dose of SCH-23390 (0, 10, 30, 50 ug/kg) for novel (left) and familiar (right) blocks, respectively. (**B**) The effect of D2 receptor antagonism (haloperidol: 0, 5, 10 ug/kg) on WSLS behavior. Plotted in same manner as (A). Asterisk indicates main effect of drug dose (*p < 0.05, 2-way ANOVA). (**C**) Regional homogeneity (ReHo) analysis of resting-state fMRI with pharmacological dopamine receptor manipulation. The clusters with significant ReHo values (p < 0.05, cluster-corrected) are superimposed on a coronal image from a standard anatomical template. (**D**) The effect of dopamine receptor antagonists on ReHo value. Bar graph indicates average and SEM of ReHo value of the voxels in the vlPFC ROI for each drug condition with individual data points superimposed (*p < 0.05, 1-way ANOVA).

Finally, to probe whether dopamine receptor-mediated manipulation of behavioral strategies also influences neural activity in vlPFC, we performed a resting-state fMRI experiment with the same dopamine antagonists (N = 7, see **Table 1**). In our previous study using the same dataset we reported that the D1 and D2 receptor manipulation induced brain-wide functional connectivity changes, most notably in the cortico-cortical and fronto-striatal FCs^28^. Here, we specifically focused on neural activity in the vlPFC region by analyzing regional homogeneity (ReHo). We chose this analysis approach as ReHo is sensitive to local changes in neural activity^29, 30^. The dopaminergic drugs induced different patterns of activity changes (ReHo signal) in bilateral vlPFC during resting-state neuroimaging (**Fig. 6C**). D2 receptor antagonist haloperidol, but not D1 receptor antagonist SCH-23390 or saline, increased vlPFC activity (1-way repeated-measures ANOVA, main effect of drug, F_(2,90)_ = 3.4, p = 0.036) (**Fig. 6D**). While these data with dopaminergic manipulation were not acquired in the context of task performance, our results suggest that the WSLS strategy that is associated with vlPFC activity is dependent on the function of dopamine D2 receptors, and that pharmacological manipulations of dopamine via D2 receptors has a clear impact on the brain circuit that vlPFC is embedded within.

## Discussion

We examined the brain-wide mechanisms underlying decision-making in different learning contexts in macaques. In a probabilistic learning task animals’ behavior was influenced by the learning context and by the preceding reward outcome. Specifically, animals used win-stay/lose-shift strategies to different degrees depending on whether they were learning new stimulus reward associations or exploiting known associations. When we analyzed the brain-wide neural activity, vlPFC stood out as a key region where both behavioral strategies and reward outcomes were encoded. Specifically, vlPFC encoded behavioral strategies during novel learning contexts. Functional connectivity in the pathways between vlPFC-ACC and vlPFC-MD was related to distinct aspects of the animals’ decisions that were dependent on the learning context. Pharmacological experiments further revealed that the manipulation of dopamine D2 receptors influenced monkeys’ behavioral strategy during learning as well as vlPFC neural activity at rest. Taken together, our results suggest a critical role of vlPFC and its associated neural networks in adaptive behavior during probabilistic learning.

The vlPFC has long been implicated in higher-cognitive function, however, the precise role of this area in learning and decision-making has only recently come into focus. Early lesion studies highlighted that damage to this area leads to a deficit in the implementation of high-order decision-making strategy in non-human primates, especially lose-shift strategies^13, 31, 32^. More recent studies using chronic or transient lesions have demonstrated a causal link between the function of this region with associative learning processes in probabilistic settings where the history of reinforcement has to be used^7, 8, 10, 12^. This prior work did not, however, directly compare the role of vlPFC across multiple learning contexts. To address this question, we designed a paradigm where subjects made choices between novel or familiar stimuli in separate blocks of trials and analyzed the pattern of choices as well as whole-brain neural activity across different learning contexts. Our behavioral analysis showed that the animals employed distinct behavioral strategies depending on the context that they were in. In the blocks of familiar trials, WSLS strategies based on monitoring of the preceding reward outcome and altering behavior accordingly were more prominent^19, 20^, whereas in the novel context the use of WSLS gradually increased as learning progressed. By comparing whole-brain fMRI signal between novel and familiar contexts, we found that vlPFC activity encoded reward outcome in both contexts within a similar time course, while the same area encoded behavioral strategy preferentially in novel contexts. This apparent disconnect between behavior and neural activity within vlPFC is notable and may relate to the fact that the vlPFC is contributing to both the learning of stimulus-reward associations and behavioral strategies in the novel context.

Notably, our vlPFC cluster that co-encoded reward outcome and ongoing learning context was mainly localized to the ventral surface of the frontal cortex, within the areas 12o/l but also extending into the anterior part of agranular insular cortex^33^ (**Fig. 2**). The location of these activations was similar to the areas previously reported in neuroimaging studies using associative learning tasks in macaques^7, 10^. This notion is also consistent with recent neural recording studies that showed a substantial reward probability or uncertainty coding in the ventral frontal cortex^9, 11, 14^, and a recent analysis of functional interactions showing a specific role for inputs from agranular insula to area 12o during feedback processing^34^. Taken together, our study reveals a new role for these parts of the ventral frontal cortex in adjusting behavior in uncertain environments.

Beyond the vlPFC itself, we found that activity in this part of frontal cortex varied with other parts of the brain during the different learning contexts. The FC between vlPFC and ACC tended to encode WSLS behavior when outcomes were delivered in the novel but not familiar blocks of trials. Further, the greatest difference between the connectivity in this pathway between novel and familiar blocks occurred when monkeys decided to switch to a different option after they failed to receive a reward (**Fig. 4B**). Notably, the activity in this pathway was marginally related to better performance, indicating that dynamic interaction between vlPFC and ACC to guide lose-shift strategies is potentially related to better behavioral performance. A specific role for this pathway in changing behaviors after failing to receive a reward agrees with reports that aspiration lesions of vlPFC result in a failure to use lose-shift strategies when learning novel associations^32^. Further, a number of prior investigations have highlighted a role for ACC in driving animals to switch to alternative options that are thought to be of higher value^21, 35^. Our data suggests that interaction between vlPFC and ACC is essential for guiding choices when the value of the perceived best option drops to a point where it is below the opportunity cost of changing behaviors.

In contrast to the role of vlPFC-ACC interactions, FC between vlPFC and MD thalamus increased when the subject decided to repeat their choice of a particular stimulus (‘stay’) even when the preceding trial wasn’t rewarded in the novel context. Such a pattern suggests that this connection encodes choice *per se* rather than a strategy to facilitate learning performance. MD has been implicated in probabilistic learning^23, 24, 36^, and prior reports from lesion studies in macaques have highlighted that MD is essential for promoting decision to stay with a particular course of action during learning^37^. Thus, our finding that there was higher functional connectivity between vlPFC and MD on stay trials appears to indicate that such lesion effects are in part caused by disconnecting this area from vlPFC.

Our results highlight a set of circuits centered on vlPFC that coordinate the flexible adjustment of behaviors in different learning contexts. It is reasonable to ask how and where the information regarding outcome and learning context converge and transform into a behavioral strategy that leads to a decision; addressing this question will require additional experiments using paired neurophysiology recordings and/or causal interrogation of specific neural circuits using viral techniques^38^.

We found that systemic administration of D2 antagonist haloperidol increased the use of WSLS strategy exclusively in novel blocks, while D1 antagonist SCH-23390 did not (**Fig. 6**). Such a pattern of effects suggests a direct link between dopamine function via D2 receptors and the learning context-dependent behavioral strategy. This notion is in line with previous literature that has emphasized the role of dopamine in a wide variety of frontal-related cognitive functions, such as working memory, motivation, attention, and learning^26, 27, 39^. Specifically, a recent study demonstrated a critical role for the meso-vlPFC dopaminergic pathway in probabilistic decision-making^14^, suggesting that dopaminergic inputs modulate vlPFC-centered functional circuits. Our resting-state fMRI data analysis revealed that the administration of haloperidol, but not SCH-23390, enhanced regional activity specifically within vlPFC. Thus, both dopaminergic modulation via D2 receptors and probabilistic learning/use of behavioral strategies appear to generally activate vlPFC. Taken together, our series of experiments reveal that a network of areas centered on vlPFC and including dopaminergic mechanisms underlie the flexible adjustment of behavioral strategy depending on the learning context.

A number of psychiatric conditions are characterized by maladaptive behaviors in uncertain reward environments. Previous studies showed that the ability to associate stimuli to probabilistic reward outcome or flexibly adapt risk tolerance in probabilistic paradigms was impaired in human or non-human primate subjects with damage to prefrontal cortex^13, 40, 41^ or patients with gambling disorders^4, 42–44^. Interestingly, a recent study showed that lesions of MD thalamus, the area we found to interact with vlPFC during decisions to stay, are associated with aberrant switching choices akin to the behavioral patterns of human subjects with paranoia^37^. This potentially implicates dysfunction within this pathway as contributing to delusional beliefs in disorders like schizophrenia. It is also noteworthy that dopamine function, particularly through D2 receptors, has been implicated in schizophrenia, the behavioral pattern of which is also characterized by impairment in flexible decision making based on probabilistic associations^5, 6, 45, 46^. Indeed, current theory posits that aberrant interactions between the salience and fronto-parietal networks, which include vlPFC as a main hub^16^, potentially underlie the biases in decision making that are observed in schizophrenia^47, 48^. Thus, the present study provides a new insight regarding the neural mechanisms underlying the flexible adjustment of behavioral strategy depending on the learning context that is potentially relevant to psychiatric disorders with impaired cognitive flexibility.

## Methods

### Subjects

Eight rhesus macaques (*Macaca mulatta,* 7-8 years old, 5 females) served as subjects. A subset of four monkeys (monkeys Ee, Ge, Me, St) underwent awake-fMRI scans. Another subset of seven monkeys (monkeys Bu, Cy, Ee, Me, Pi, St, Wo) underwent pharmacological experiments with dopaminergic drugs. The experiments performed for each subject are summarized in **Table 1**. All procedures were reviewed and approved by the Icahn School of Medicine Animal Care and Use Committee.

### Surgery

Prior to training, an MRI compatible head-fixation device (Rogue research, Cambridge, MA) was surgically implanted using dental acrylic (Lang Dental, Wheeling, IL) and ceramic screws (Thomas Research Products, Elgin, IL) in the animals that underwent behavioral testing (monkeys Ee, Ge, Me, Pi, St). Briefly, following induction with ketamine (5 m/kg) and dexmedetomidine (0.0125 mg/kg), the animals were maintained on isoflurane (2-3%), and 8-10 screws were implanted into the cranium and the head fixation device was bonded to the screws using dental acrylic. The animals were treated for discomfort and monitored by the researchers and veterinary staff till fully recovered. The position of implant was determined based on a pre-acquired T1-weighted MR image.

### Probabilistic learning task

We used a reward-based probabilistic learning task that we recently developed for macaque monkeys^28^. The task was controlled by NIMH MonkeyLogic software^49^ running on MATLAB 2019a (MathWorks, Natick, MA) and presented on a monitor in front of the monkey. In this task, animals were required to choose between two visual stimuli using a directed saccadic eye movement. A trial began with the appearance of a fixation spot (white cross) at the center of the screen, which the monkey had to maintain fixation on for 1-1.5 sec to initiate a trial. The fixation spot was then extinguished and two stimuli were simultaneously presented to the right and left on the screen. The two stimuli presented on each trial were randomly chosen from a larger pool of three visual stimuli that were associated with different reward probabilities (0.9, 0.5, and 0.3). Each trial fell into one of three categories based on the reward probabilities of the options presented: High-Low (0.9-0.3), High-Mid (0.9-0.5), and Mid-Low (0.5-0.3). Stimuli were either novel at the beginning of each block of 100 completed trials (novel block), or subjects had previously learned the probability of receiving a reward associated with each image, making them highly familiar (familiar block). Once stimuli were presented, subjects were required to move their gaze toward either the right or left stimulus option (‘response’) within 2 seconds. Following a response, the chosen stimulus remained on screen for 0.3 sec, and then was removed and a reward (1 drop of apple juice) was provided in accordance with the probability of the chosen option. Subsequently an inter-trial interval (ITI, 3-3.5 sec) followed. A trial with a fixation break during the fixation period or with no response within the response window was aborted; all stimuli were extinguished immediately and ITI started. The same trial was repeated following an aborted trial.

The animals were trained in a mock MRI scanner for 3-6 months in advance of experiments. On an experimental day, the animals performed 2-6 blocks, in which novel and familiar blocks (i.e. learning context) were pseudorandomly interleaved. In awake-fMRI sessions, animals performed the task in the MRI scanner during functional scans (see *Awake fMRI data acquisition* section). In pharmacology sessions, animals performed the task in the mock scanner. The injection of saline, SCH-23390, or haloperidol solution (i.m.) was performed 15 minutes prior to the task start, and the order of drug treatment was randomized. The injections were at least a day apart (SCH-23390) or a week apart (haloperidol) to avoid potential prolonged effects of the drug, in accordance with known pharmacokinetics of the drugs in macaque monkeys^50^.

### Awake fMRI data acquisition

Animals sat in sphynx position in a custom-built MRI-compatible primate chair (Rogue Research, Cambridge, MA) to perform a behavioral task in the MRI scanner (Siemens Skyra 3T). First the animals received an intravenous injection of a contrast agent, monocrystalline iron oxide nanoparticle or MION (BIOPAL, Worcester, MA), at a concentration of 10 mg/kg 30 minutes prior to the scan^51, 52^. After head fixation, a custom-built 4-channel coil was placed around the head. Eye movement was monitored via infra-red camera and tracked using EyeLink 1000 software (SR Research, Ottawa, Canada). Juice reward was provided through pressurized tubing. A session started with a set of setup scans which included shimming based on the acquired fieldmap. Following a 3D T1-weighted MPRAGE image (0.5 mm isotropic, TR/TI/TE 2500/1200/2.81 ms, flip angle 8°), 2-6 runs of echo planar image (EPI) functional scans (1.6 mm isotropic, TR/TE 2120/16 ms, flip angle 45°, 300-500 volumes per each run) were obtained, with each functional scan occurring in conjunction with a separate block of behavioral testing. Overall, animals each completed 4 to 7 scanning sessions, for a total of 23 scanning sessions (55 novel and 42 familiar blocks).

### Resting-state fMRI data acquisition

The scans were performed under the same protocol we previously developed for macaque monkeys^28, 53, 54^. In brief, following sedation with ketamine (5mg/kg) and dexmedetomidine (0.0125mg/kg) the animals were intubated. They were then administered MION (10 mg/kg, i.v.), and three EPI functional scans (300 volumes per each run) were obtained, along with a T1-weighted structural scan (pre-injection scans). Following drug i.v. injection (saline, SCH-23390, or haloperidol) and 15 minutes waiting period, another set of three functional scans was acquired (post-injection scans). Low-level isoflurane (0.7-0.9%) was used to maintain sedation through a session so that neural activity was preserved while minimizing motion artifacts. The doses of drugs used in the scans (50 µg/kg and 10 µg/kg for SCH and haloperidol, respectively) were pre-determined based on a prior PET study to achieve up to 70-80% occupancy of the DA receptors in macaques^50^. Vital signs (end-tidal CO_2_, body temperature, blood pressure, capnograph) were continuously monitored and maintained as steadily as possible throughout an experimental session.

### Drugs

SCH-23390 hydrochloride (Tocris Bioscience, Minneapolis, MN) and haloperidol (Sigma-Aldrich, St. Louis, MO) were used as D1 and D2 receptor selective antagonists, respectively^55^. Both SCH and haloperidol were dissolved and diluted in 0.9% saline to achieve target dose of 1 ml solution. 0.9% saline (1 ml) was also used as a control solution. The solution was prepared fresh on every experimental day.

### Behavioral data analyses

All behavioral data was analyzed using MATLAB 2022b. Choice performance was defined as the proportion of trials in a block (100 trials) in which monkeys chose an option associated with higher reward probability in the stimulus pair presented. Reaction time (RT) was defined as the duration from the timing of visual stimuli presentation to the timing of response initiation. Choice performance was computed for bins of 10 trials at each block and averaged for each subject, then finally averaged across subjects for each context, novel and familiar. We also computed choice performance for each quartile and performed a two-way repeated-measures ANOVA (trial bin: 1-4, block type: novel or familiar) for each block type. We reasoned that a significant interaction of trial bin by block type (p < 0.05) indicates the improvement of performance through successful learning in novel blocks. Choice performance and RT on each stimulus pair were assessed by two-way repeated-measures ANOVA (stimulus pair: 0.9-0.3, 0.9-0.5, 0.5-0.3, block type: novel or familiar).

Switching trials were defined as trials in which the monkeys chose a different stimulus although the previously-chosen stimulus was available in the current trial, as opposed to stay trials in which the monkeys chose the same stimulus sequentially. Thus the proportion of switching trials was limited to those trials in which the previously-chosen stimulus was available. The proportion of switching trials regarding previous outcome and block type was assessed using two-way repeated-measures ANOVA (outcome: reward or no-reward, block type: novel or familiar). We interpreted a significant interaction of outcome by block type (p < 0.05) to indicate that the proportion of win-stay/lose-shift (WSLS) trials was varied depending on the learning context. We also performed a two-way repeated-measures ANOVA (trial bin: 1-4, block type: novel or familiar) on the proportion of WSLS choices to assess the impact of learning on WSLS strategy. The direct relationship between WSLS choices and choice performance was assessed by calculating the Pearson’s correlation coefficient. To do this with all subjects combined, the proportion of WSLS choices and the choice performance on each block were z-transformed for each session and each block type.

The effect of drug injection on the proportion of WSLS choices was assessed by two-way repeated-measures ANOVA (trial bin: 1-4, dose: 0, 5, or 10 µg/kg of haloperidol, or 0, 10, 30, or 50 µg/kg of SCH-23390) for each block type. The significant main effect of drug dose (p < 0.05) indicates the impact of the drug on the proportion of WSLS trials in a specific learning context.

### fMRI data analyses

Imaging data were analyzed using a customized AFNI processing pipeline for non-human primates^56^ and the standard NMT atlas^57^. Following the preprocessing steps, whole-brain analysis was performed. For awake-fMRI data, regression analysis was performed for each session with the timing of the outcome as a regressor. One scanning session with only novel blocks was excluded from this analysis, resulting in 22 sessions (51 novel and 42 familiar blocks). The resulting correlation coefficients for each voxel were submitted to a two-way ANOVA (outcome: reward or no reward, block type: novel or familiar) with subject and session as random effects^10^. Group-level statistics were computed by 3dClustSim using initial thresholding at p < 0.05 in the ANOVA and a cluster size of 14 voxels that is corrected for multiple comparisons at p < 0.05. Subsequent conjunction analyses specified the areas that survived cluster-based correction in both outcome and block type (context) contrast.

A region of interest (ROI) was chosen based on the result of the conjunction analysis. The peak voxel from right vlPFC and its adjacent voxels (faces touching) was used as the main ROI for subsequent ROI-based analyses. First, the time series was extracted from the ROI and z-transformed for each block, and the timing of the signal was aligned to the timing of outcome. Then, trials were divided based on outcome (reward or no reward) and subsequent decision (stay or shift), and the averaged time series for each trial type with smoothing was computed to create a peri-stimulus timing histogram for each task. A sliding-window multiple linear regression (500 msec window, 100 msec step) concerning outcome and stay/shift decision was performed and the resulting beta coefficients were plotted around the timing of outcome. Also, a correlation between ROI value (0-4 sec from the timing of outcome) and normalized choice performance was calculated. A multidimensional analysis was performed by plotting beta coefficients (outcome and stay/shift decision coding) in 2-D space for each block type separately. Then, the Euclidean distance between novel and familiar blocks across the timing of the trial was computed and plotted against the trial time-course. The distance measure was compared to shuffled data (95% CI) that was computed by randomly assigning block types for each trial and iterated 1,000 times.

Subsequently, functional connectivity (FC) analysis was performed using the right vlPFC seed that was used in the main ROI analysis. We performed a generalized form of context-dependent psychophysiological interactions (gPPI)^22^ to compute a vlPFC-derived network modulated by learning context (novel or familiar) at the whole-brain level. Then, a secondary ROI was defined based on peak voxels in the brain map. The ROI time-course of FC was computed by calculating the seed-ROI correlation (Pearson’s r) over time using a sliding window analysis. The trials were divided based on outcome and subsequent choice, and the FC time course for each trial type was calculated around the timing of outcome, for each block type. The significant FC change between -4 to 8 seconds after the outcome timing was detected when 3 consecutive bins reached p < 0.05 in a Wilcoxon’s rank-sum test. We further performed a sliding-window multiple regression analysis concerning outcome and stay/shift decision in the FC. The beta coefficients of the interaction of outcome by stay/shift decision were plotted as WSLS coding of the FC and compared to the 95% CI of the shuffled data.

Resting-state fMRI data were preprocessed in the same manner as the awake scans. The residual error file for each run was split in half, resulting in 6 runs for pre- and post-injection scans, respectively, for each of the drug injection sessions. Regional homogeneity (ReHo) analysis was performed using the function 3dReHO in the AFNI FATCAT toolbox^30^. ROI analysis was performed using the right vlPFC ROI that was used in the main ROI analyses. Two-way ANOVA (drug: saline, SCH-23390, or haloperidol, injection: pre or post) was performed to assess the effect of drug injection on vlPFC activity. The results were superimposed on the NMTv2.0 template for visualization purpose^57^.

## Acknowledgments

A.F., C.E., B.E.R., and P.H.R. are supported by grants from the BRAIN initiative (R01MH117040). B.E.R. is supported by grants from NIMH (R01MH111439) and NINDS (R01NS109498). A.F. is supported by Overseas Research Fellowship from Takeda Science Foundation and a Brain & Behavior Research Foundation Young Investigator grant (#28979). We would like to thank Dr. Paula Croxson for providing the foundation on which this work was built, Dr. Yukiko Hori for advising on drug preparation, and Jairo Munoz and Niranjana Bienkowska for assistance with data acquisition. For help with fMRI data pre-processing and analysis we thank Drs Paul Taylor, Alex Franco, and Vincent Costa.

## Author contributions

A.F., C.E., B.E.R., and P.H.R. designed the study. A.F., C.E., S.H.F., and L.F. performed the study. A.F. analyzed the data. A.F., C.E., B.E.R., and P.H.R. wrote the original draft. All authors edited the paper.

## Data Availability

The data that support the findings of this study are available from the corresponding authors upon reasonable request.

## Supplementary Information for

**Supplementary Figure 1.**
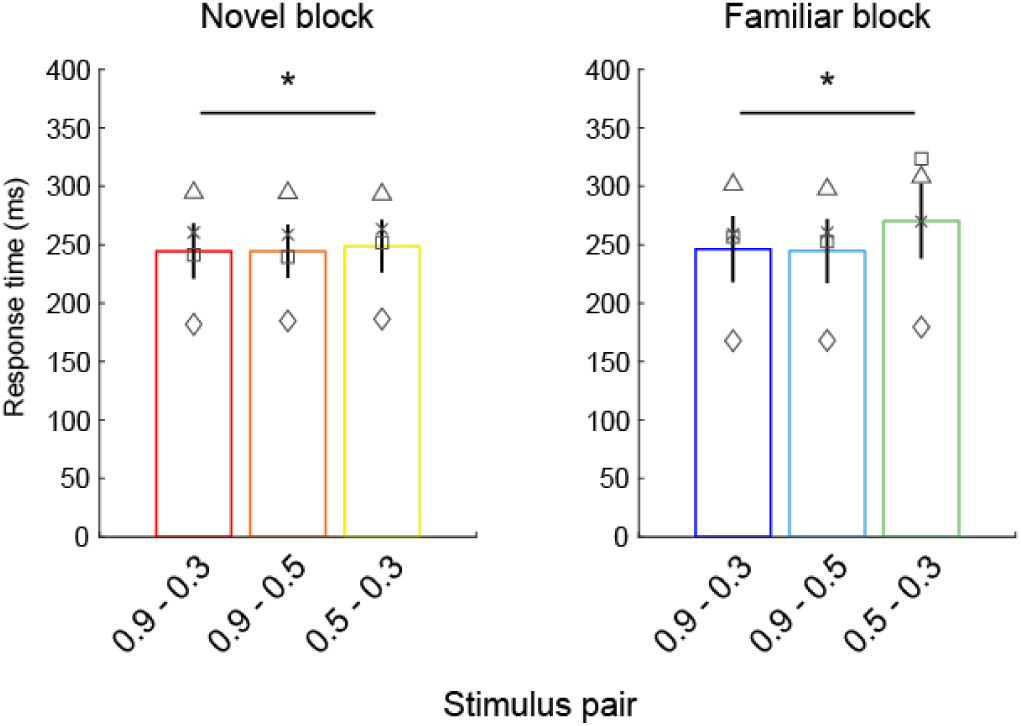
Response time (RT) of monkeys. Bar graphs show average and SEM of RT for each stimulus pair in novel (left) and familiar (right) blocks. Symbols represent each animal. Asterisks indicate significant main effect of stimulus pair (*p<0.05, 2-way repeated-measures ANOVA).

**Supplementary Figure 2.**
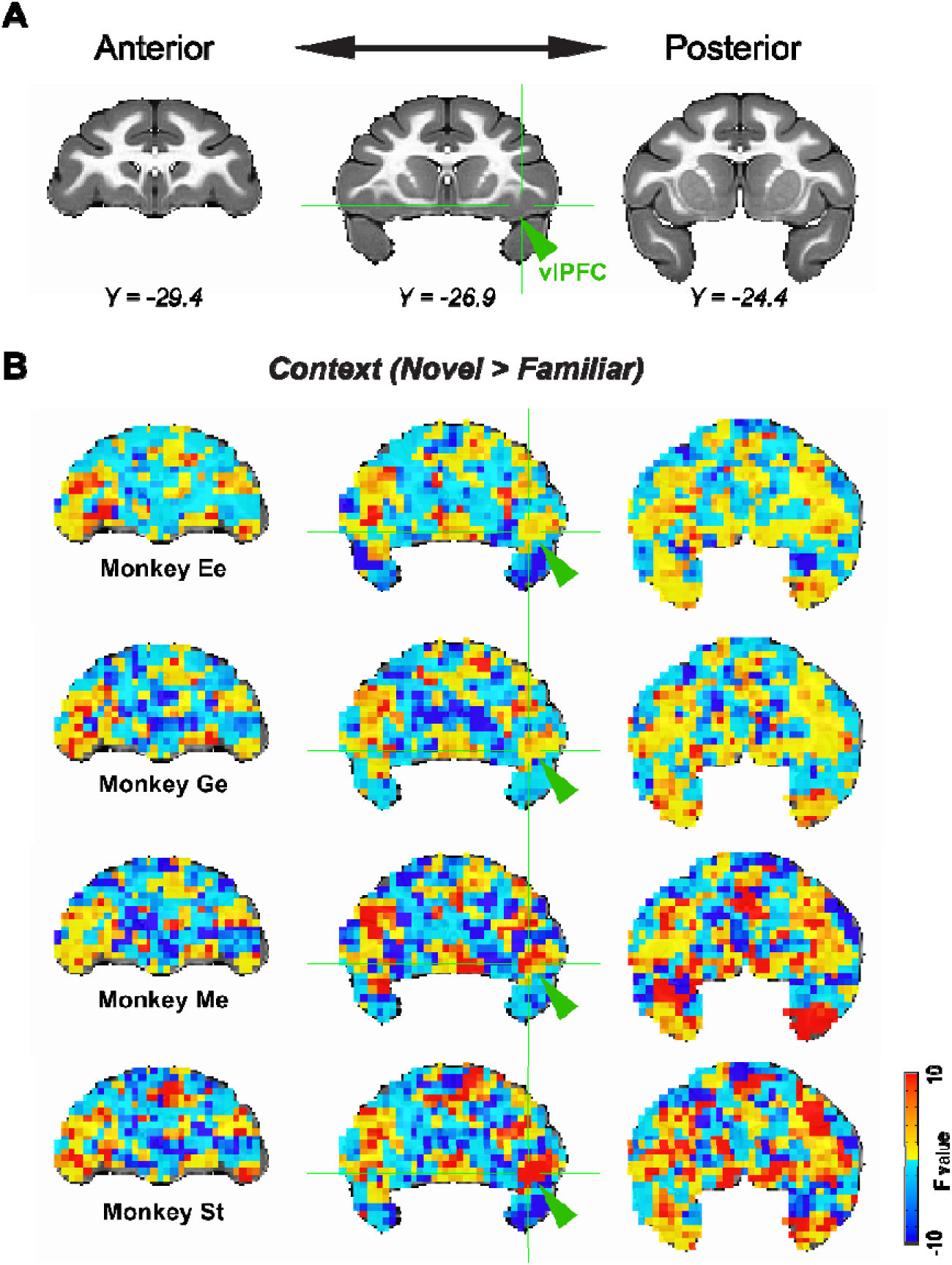
Context coding in individual monkeys. (**A**) Anatomical templates showing coronal slices around vlPFC ROI. (**B**) Unthresholded map of F-stats superimposed on the anatomical templates in (A). The data for each animal is shown in each row. Crosshairs and arrowheads indicate the peak coordinates of vlPFC ROI used in time-course analyses.

**Supplementary Figure 3.**
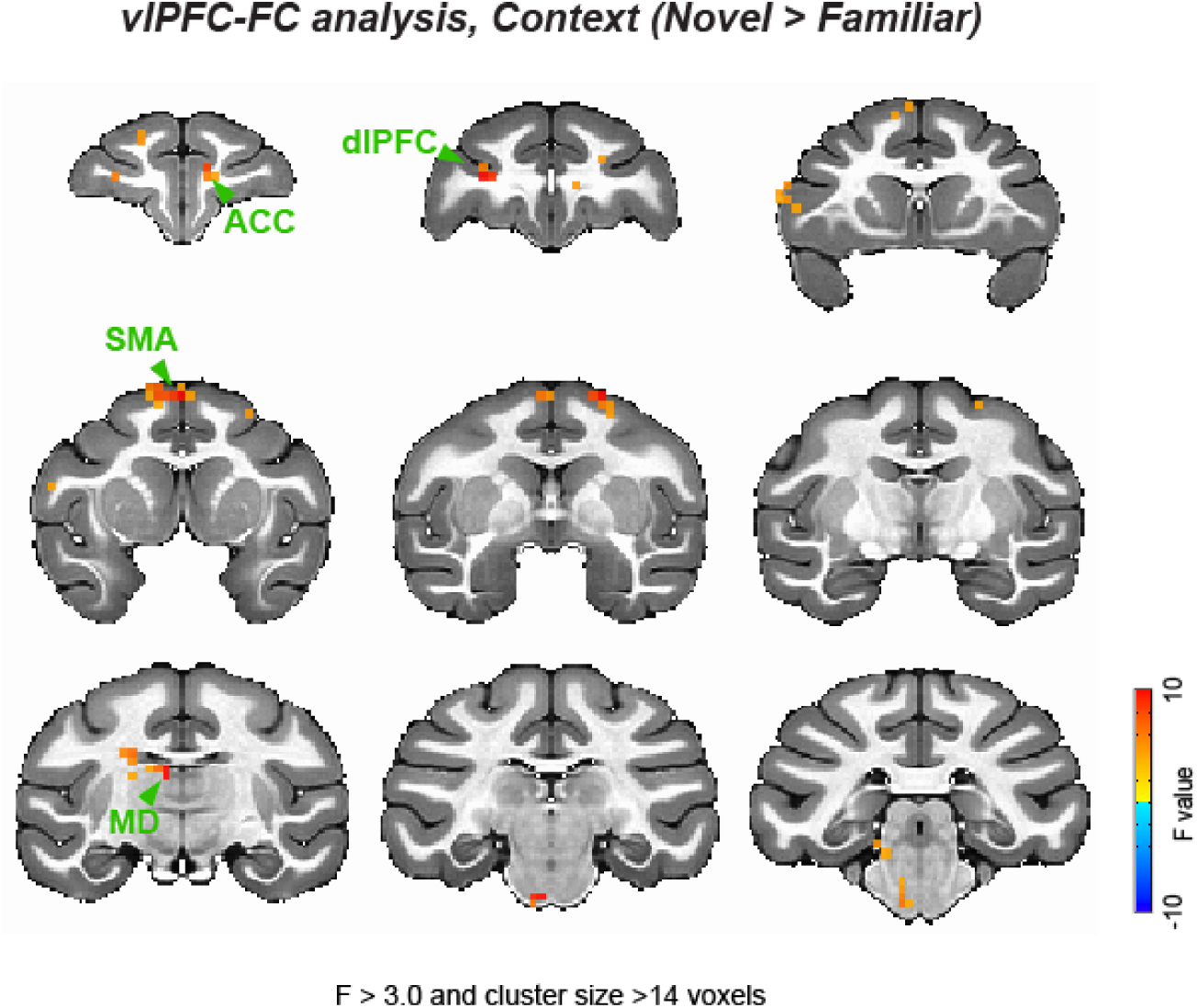
Functional connectivity analysis using vlPFC seed. Whole-brain map of F-stats in significant clusters (p < 0.05, cluster-corrected, generalized psycho-physiological interaction or gPPI) superimposed onto an anatomical template. Coronal slices (4.0 mm apart) are shown from anterior (top left) to posterior (bottom right) planes.

**Supplementary Figure 4.**
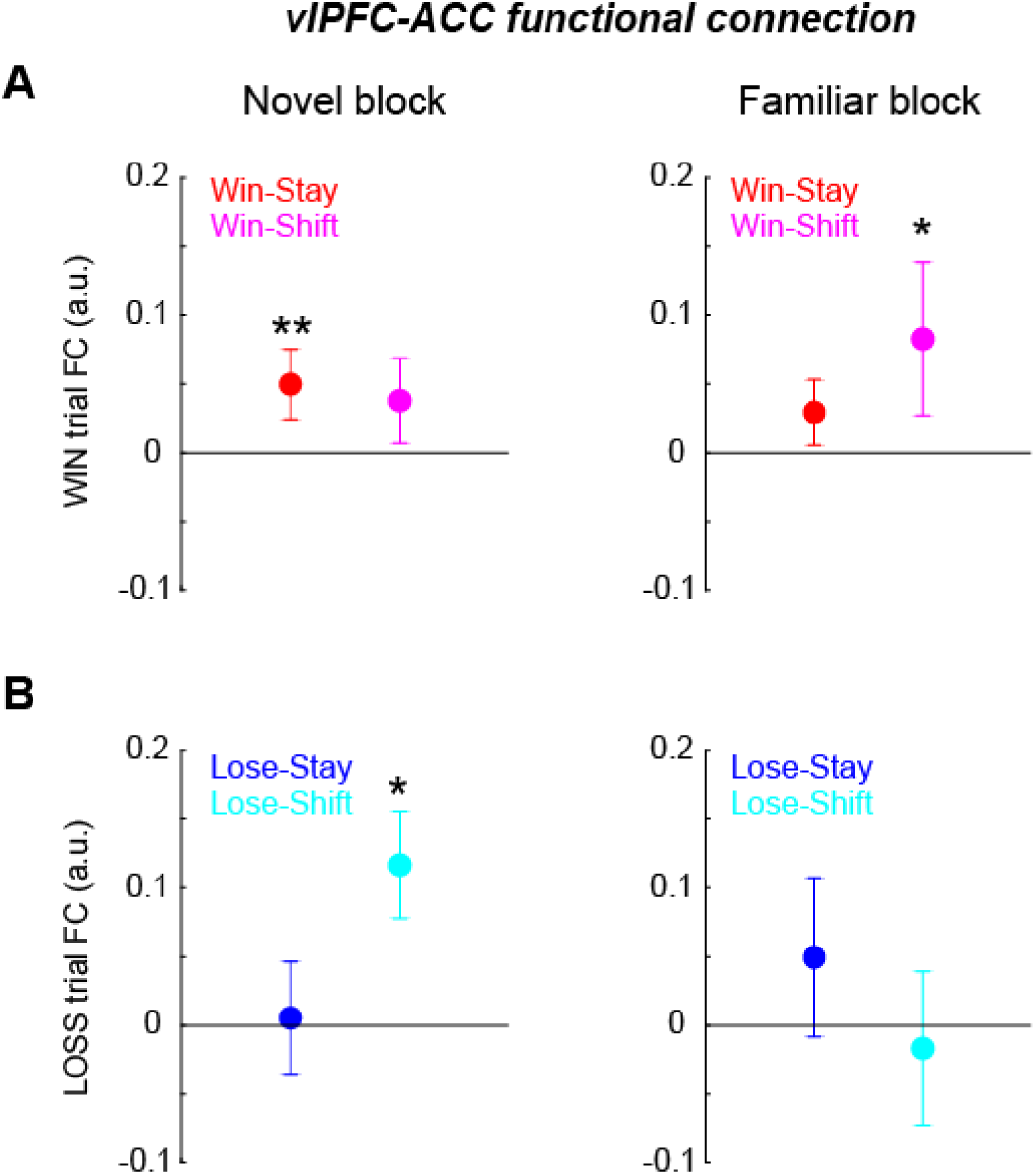
The impact of outcome and stay/shift decision on vlPFC-ACC functional connection. The average FC between vlPFC and ACC around the timing of outcome (-2 to +2 seconds after outcome) are plotted for win-stay and win-shift trials (A) and lose-stay and lose-shift trials (B) for novel (left) and familiar (right) blocks, respectively. Error bar indicate SEM. Asterisks indicate significant FC changes from zero (**p < 0.01 or *p < 0.05, rank-sum test).

**Supplementary Figure 5.**
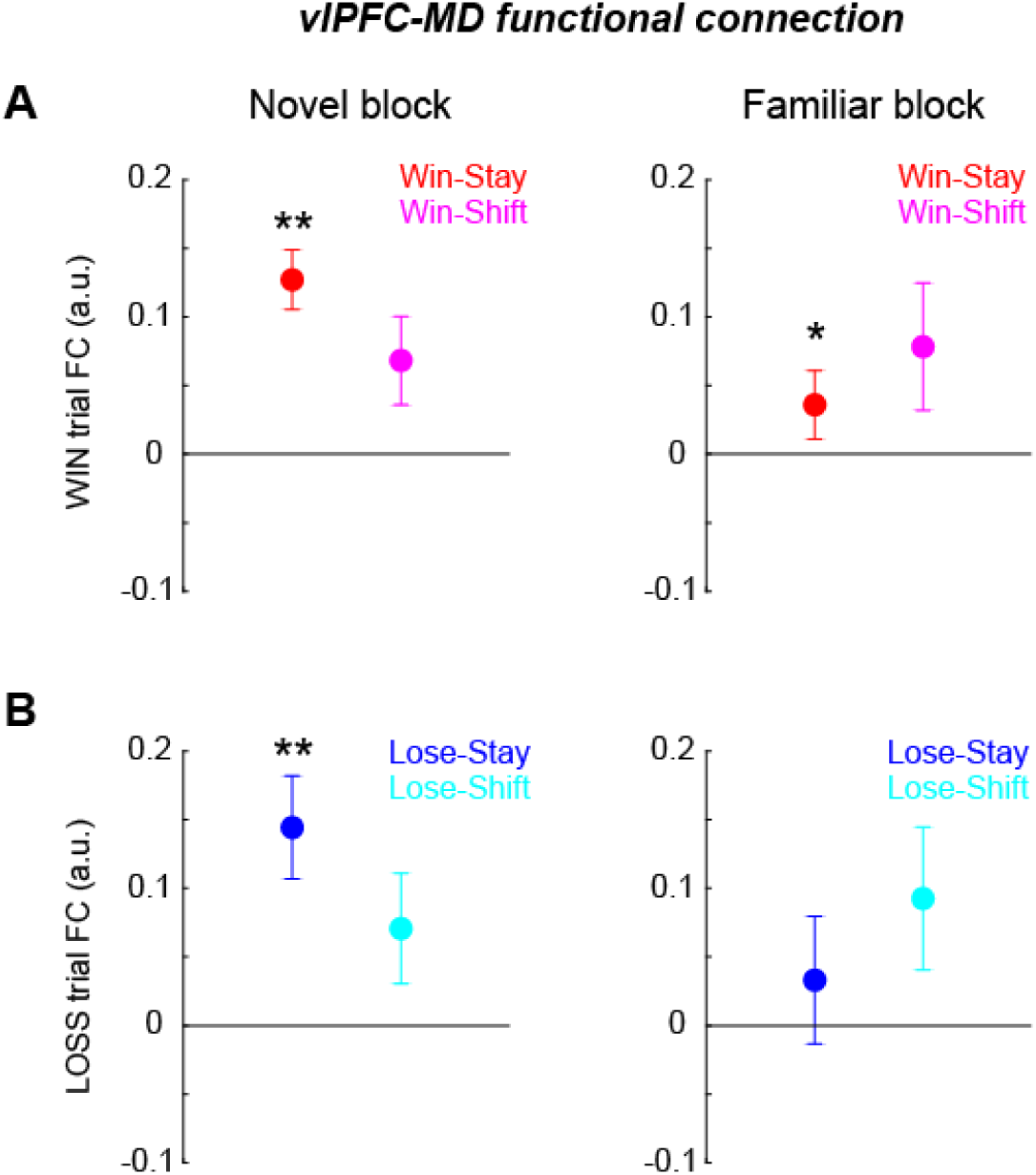
Functional connection between vlPFC and MD thalamus around the outcome timing. (**A, B**) The average FC between vlPFC and MD around the timing of outcome are plotted for novel (left) and familiar (right) blocks, respectively. The conventions are the same as Figure S4. Asterisks indicate significant FC changes from zero (**p < 0.01 or *p < 0.05, rank-sum test).

**Supplementary Table 1.**
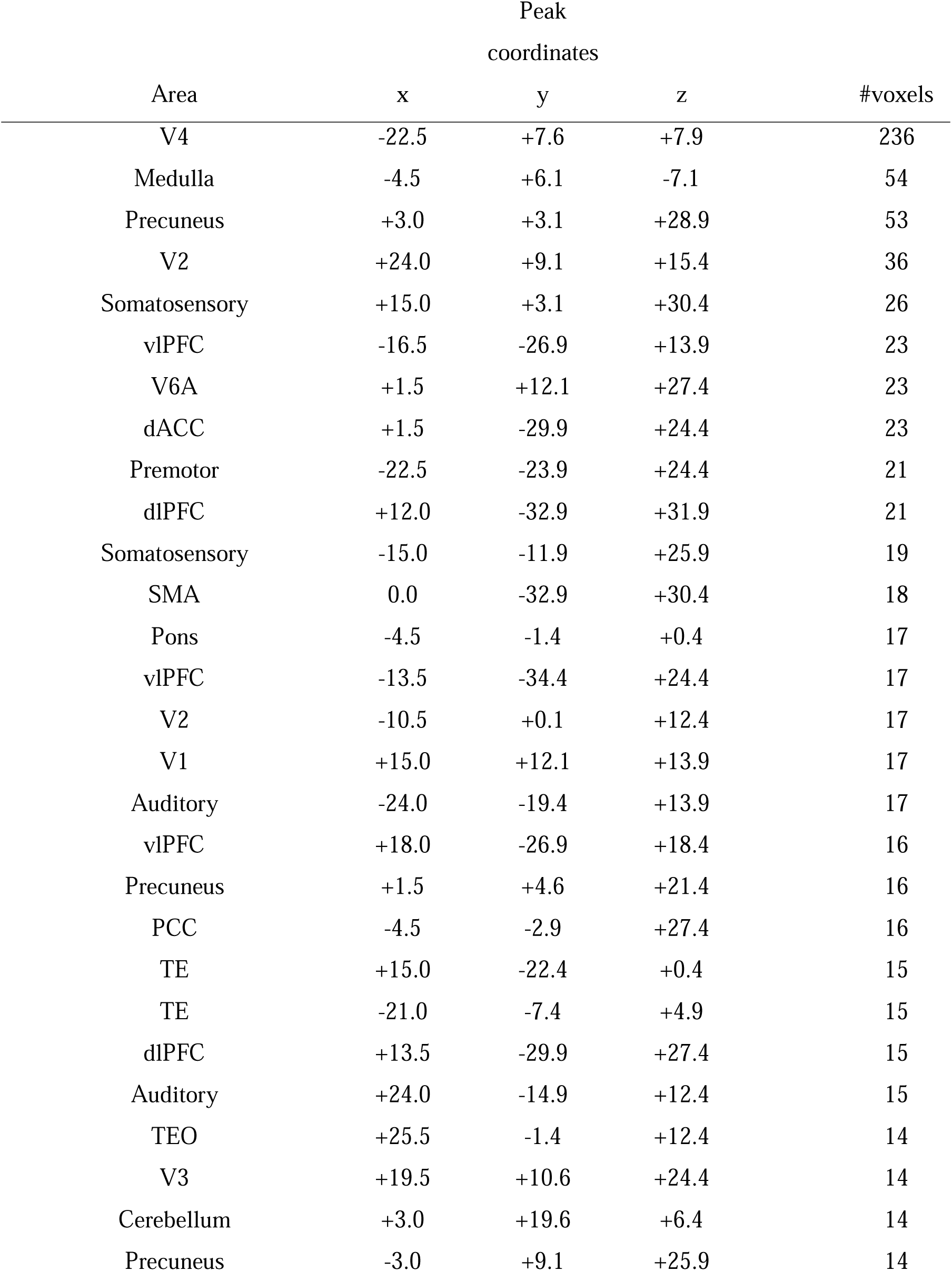
Full list of clusters in the whole-brain analysis that encoded learning context and reward outcome. dlPFC: dorsolateral prefrontal cortex, vlPFC: ventrolateral prefrontal cortex, V1: primary visual cortex, V2: secondary visual cortex, V3: third visual cortex, V4: fourth visual cortex, TE: anterior inferotemporal cortex, TEO: posterior inferotemporal cortex, dACC: dorsal anterior cingulate cortex, PCC: posterior cingulate cortex.

**Supplementary Table 2.** Full list of clusters in the functional connectivity (gPPI) analysis that encoded learning context in relation to right vlPFC seed timeseries. dlPFC: dorso-lateral prefrontal cortex, V1: primary visual cortex, V2: secondary visual cortex, dACC: dorsal anterior cingulate cortex.

| Area | Peak<br>coordinates |  |  | #voxels |
| --- | --- | --- | --- | --- |
|  | x | y | z |  |
| Pons | +1.5 | -7.4 | -1.1 | 63 |
| dlPFC | +7.5 | -35.9 | +31.9 | 40 |
| Somatosensory | +6.0 | +7.6 | +34.9 | 37 |
| Premotor | 0.0 | -22.4 | +36.4 | 29 |
| Premotor | -9.0 | -19.4 | +36.4 | 27 |
| Cerebellum | -3.0 | +4.6 | +1.9 | 25 |
| V2 | -3.0 | +10.6 | +19.9 | 24 |
| MD thalamus | +3.0 | +10.4 | +19.9 | 22 |
| V1 | +7.5 | +19.6 | +16.9 | 21 |
| Medulla | +1.5 | +1.6 | -7.1 | 19 |
| V1 | -9.0 | +16.6 | +28.9 | 19 |
| V2 | +9.0 | +13.6 | +24.4 | 17 |
| Premotor | +21.0 | -23.9 | +21.4 | 15 |
| V1 | -12.0 | +19.6 | +4.9 | 14 |
| dACC | -4.5 | -34.4 | +25.9 | 14 |

